# Ants hold a grudge: Associative learning of non-nestmate cues improves enemy recognition

**DOI:** 10.1101/2023.12.19.572127

**Authors:** Mélanie Bey, Rebecca Endermann, Christina Raudies, Jonas Steinle, Volker Nehring

## Abstract

Recognition protects biological systems of all scales, from cells to societies. Social insects recognize their nestmates by colony-specific chemical labels, which individuals store as “templates” in their memories. The distributed model of nestmate recognition predicts that individual experiences cause variation between the recognition templates that different individuals store in their memory. Here, we show for the first time that ants associatively learn recognition labels of both friends and enemies during brief encounters. Because different individuals will accumulate different experiences throughout their lives, their later decisions on whether to accept or reject other individuals will also vary. Individual associative learning can thus explain many phenomena from nasty neighbour effects to age polyethism. To avoid chaos caused by inconsistent decisions across individuals, decisions are made in a distributed manner at the colony level. Group-level decision can thus be much more accurate than any individual decision alone.

## Introduction

A system consists of interrelated elements that form a unified whole. The individual elements often interact smoothly until an outside disturbance occurs, such as sand being strewn into the gears. For a system’s performance it is thus important to protect its integrity. This can be done by encasing the system in a protective layer, or, as it is often the case in biological systems that cannot be completely sealed off, through recognition. Recognition has been studied intensively at different scales from immune systems to societies. Colonies of eusocial insects are impressive systems that have allowed many species to prosper for over 100 million years^1^. For colonies to last, individuals must selectively cooperate with relatives. Nestmate recognition is the sine qua non condition that makes this possible^2,3,4^.

In ants, recognition is based on the identification of colony-specific odours (labels), which workers compare to a neural representation of their own olfactory identity (template^5^). Throughout an ant’s life, learning continuously shapes the nestmate template to keep up with the constant changes in colonial odours^6,7,8,9,10^. The main mechanism allowing template update is often thought to be that constant exposure to an odour causes ants to become less aggressive toward it^11,12^. This process is called habituation: Individuals ‘get used’ to these odours and stop reacting to them. Indeed, when ants are experimentally exposed to non-nestmate odour in a peaceful setting, they subsequently treat the former non-nestmates as if they were nestmates^13,14,15,11,16^.

Multiple studies have shown that there is a tremendous individual variation in nestmate recognition abilities. For example, older workers are known to be better at recognition, which is typically attributed to temporal polyethism, i.e. a programmed task change as individuals age^17,18^. The remaining variation is sometimes attributed to individual variation in recognition cues^19^ or even considered experimental noise^20^. A current model hypothesizes that associative learning may be an additional factor contributing to the variation in nestmate recognition^21^. This would be the case if templates were not only formed by habituation, but if workers would associate the specific experience they make with other individuals with the odour labels these others bear. When being fed or groomed by a nestmate, this ‘pleasant’ experience would act as an unconditioned stimulus (US), a stimulus that has a valence in itself and induces a specific emotion-like state. The ants associate the US with the nestmate’s odour, which thus becomes a conditioned stimulus (CS), a stimulus that gained its valence only through the association with the US. Such an association then creates a nestmate template. Analogously, being attacked by non-nestmates might act as an US that brings the ants to learn the non-nestmate’s colony odour as a CS, forming a non-nestmate template. A specific experience is connected to the odours, so that the odours take on the valence of the experience. This contrasts with habituation where the odours simply stop to matter. The accumulation of templates gained through associative learning, or their continued refinement, would constantly improve the recognition skills of each individual. That being said, not each individual needs to be perfect at recognition and the templates may vary widely between individuals. Only the aggregate of templates gathered by many individuals can lead to precise recognition at the colony level (distributed model of nestmate recognition^21^). While associative learning of non-nestmate cues could explain many phenomena observed in colony defence, its existence has never been tested.

The major prediction of the associative learning hypothesis is that highly specific behavioural changes, i.e. an increase or decrease in aggression when reacting to a specific stimulus, can be induced by relatively short encounters with other ants. In contrast, habituation would only reduce aggression and requires relatively long exposure to odours. Circumstantial evidence is in line with this prediction: In ants, foragers are known to be more aggressive than inside workers against non-nestmates^17^. While this may be a simple consequence of programmed temporal polyethism, the foragers are also the only ants that ever encountered and thus fought with non-nestmates. The nurses would not be able to learn non-nestmate odours inside their own nest. Additionally, geographical distance between colonies has been shown to affect the dynamics of aggressive interactions between the colonies. Sometimes neighbours are relatively peaceful (dear enemy phenomenon^22^), which is thought to increase fitness by reducing the cost of regular fights against an established competitor^23,24,25^. In other species, neighbour interactions are more aggressive than interactions between distant colonies, which may be a consequence of specific combinations of territory values and costs of fights (nasty neighbour effect^26,27,28^). However, experiments specifically testing the ultimate and proximate causes and consequences of the two phenomena are still lacking.

Here, we test the hypothesis that associative learning of non-nestmate colony odours is possible, and that it can explain the nasty neighbour effect. Our results are the first to demonstrate that associative learning plays a crucial role in the formation of both nestmate and non-nestmate recognition templates. The processes involved fulfil all necessary requirements for an efficient distributed nestmate recognition system as predicted by the distributed model of nestmate recognition^21^.

## Results

First, we explored the variation in aggression between colonies of *Lasius niger* ants in an ecologically relevant context, to test for a nasty neighbour effect (Experiment 1). We found that ants from colonies that were within foraging distance were more aggressive towards each other than colonies that were further apart (see Supplement 1). We suspected that the elevated aggression resulted from repeated interactions between neighbouring colonies. To test whether the aggression can be modulated by associative learning during encounters with enemies, we set up three laboratory experiments.

We explored how previous experiences with non-nestmates alter the subsequent aggression of focal ants (Experiment 2). We tested whether a succession of fights with ants from a specific non-nestmate colony would later increase the aggression of focal ants toward that colony. During the training phase, focal ants were exposed to brief encounters of 1 minute with ants from a non-nestmate colony or with a nestmate, once a day over five days. When focal ants encountered individuals from the same (later ‘known’) non-nestmate colony once per day, the aggression of the focal ants increased over the five days (Suppl. Fig. 3.1A). In contrast, when encountering nestmates during the training phase, the focal ants remained peaceful throughout (Suppl. Fig. 3.1B). After the training phase, on day six, we set up encounters with non-nestmates from either the same colony as during the training phase (known treatment), from a different non-nestmate colony (unknown treatment), or with nestmates (experimental design in Fig. 1A). The individual experience of the focal ants during the training phase affected their aggression on day six (Fig. 1B; n = 312, factor treatment p < 0.001; Suppl. Tab. 3.1): Focal ants were most aggressive when encountering known non-nestmates, i.e. when they had previously encountered ants from the same non-nestmate colony. The aggression of ants that encountered ants from a so far unknown non-nestmate colony was lower and not different from the aggression of ants that had previously only encountered nestmates (Fig. 1B; Suppl. Tab. 3.2). This could explain the nasty neighbour effect observed in experiment 1. Focal ants that were presented with nestmates on day six were uniformly peaceful, independent of whether they were trained with nestmates or non-nestmates (n = 156, p = 0.25; Suppl. Fig. 3.2; Suppl. Tab. 3.3).

**Figure 1:**
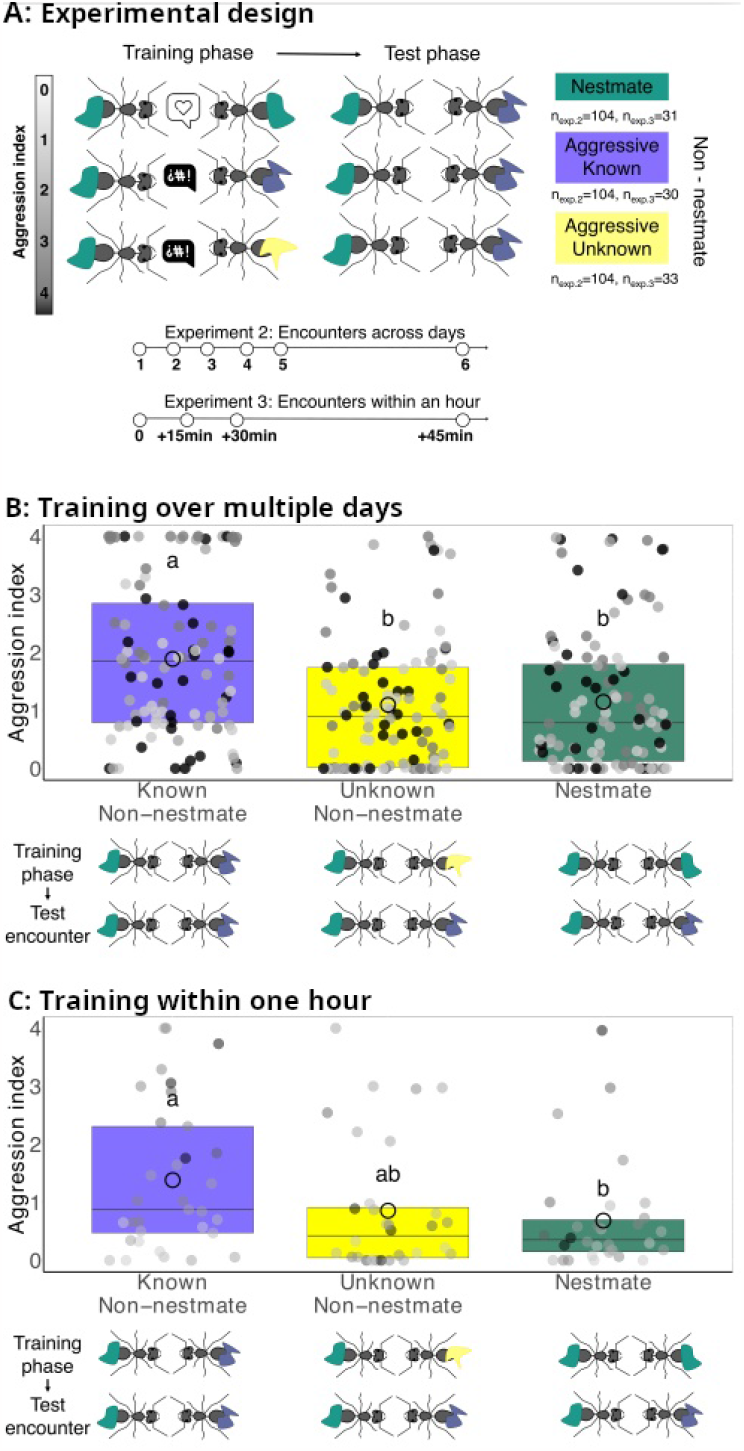
Ants are more aggressive against known than unknown non-nestmates. During a training phase, we repeatedly exposed focal ants to non-nestmates from a specific colony, or to nestmates. Later we tested whether the focal ants would attack ants from the previously encountered “known” colony, or non-nestmates from a novel, unknown colony (A). In Experiment 2 (B), we staged five training encounters, one per day. During the test on day six, focal ants were more aggressive towards known non-nestmates than towards unknown non-nestmates (n = 104 encounters per combination, p < 0.001; Suppl. Tab. 3.2). The aggression of ants that had previously only encountered nestmates was not different from that of ants encountering unknown non-nestmates (p = 0.74). In experiment 3 (C), we condensed the training into three encounters within 45 minutes. Ants still learned to recognize non-nestmates and were more aggressive if they already knew the non-nestmates than if they had previously only encountered nestmates (p < 0.05, n = 30-33 per group; Suppl. Tab. 4.2). Again, whether ants had experienced only nestmates or a different non-nestmate colony did not affect aggression (p = 0.42), but the difference in aggression between known and unknown non-nestmates was not as robust as in experiment 2 (p =0.057). For each experiment, the y-axis shows the aggression expressed by the focal ants towards non-nestmates during the test encounter. Small grey dots are measurements of individual ants’ aggression scores, the different shades correspond to the combination of focal ant and non-nestmate colony ID (2-3 different focal colonies per experiment). The large circles indicate means and the box plots medians and interquartile ranges. Groups with different letters were significantly different in a pairwise t-test (p < 0.05; Suppl. Tabs. 3.2, 4.2). The coloured shapes on the ant pictograms represent the different colony-specific odour labels. The symbols in the speech bubble indicate the tone of the encounter. <3 symbolises that the encounter was passive, !#¿ that it was agonistic.

We conducted a third experiment to evaluate whether the length of the intervals between exposures to non-nestmates affects the learning process (Experiment 3). We condensed the training phase into a single day (Fig. 1A) and only exposed the ants to non-nestmates during the test. Three learning encounters of one minute each within 45 minutes produced a similar effect as five encounters on subsequent days did (Fig. 1C, n = 94, p < 0.05, Suppl. Tab. 4.1). On the fourth encounter, focal ants were more aggressive when encountering known non-nestmates than when encountering nestmates (n = 94; Fig. 1C; p < 0.05, Suppl. Tab. 4.2). While the aggression was not significantly different between ants encountering known and unknown non-nestmates, there was a trend and the sample size was relatively low (Fig. 1C; p = 0.06, Suppl. Tab. 4.2). Compared to learning over multiple days, the effect may thus be weaker here. In this experiment, there was no evidence that aggression increased over subsequent training encounters (Suppl. Fig. 4.1).

In the fourth experiment, we tested whether being attacked constitutes an unconditioned stimulus that leads to associative learning of non-nestmate colony odours. We rendered opponent ants passive by ablating their antennae (Suppl. Fig. 5.1) and set up training encounters between the focal ants and these passive non-nestmate opponents, or aggressive non-nestmates as a control. Training happened daily for three days (Suppl. Fig. 5.2). On day four, all focal ants encountered aggressive non-nestmates, which were either from a known or unknown colony (Fig. 2A). Whether these non-nestmates had previously been aggressive predicted the aggression of focal ants on day four (Fig. 2B; n = 322; p < 0.05; Suppl. Tab. 5.1). Ants were significantly more aggressive when they encountered non-nestmates that they had previously experienced to be aggressive than if they had been passive (p < 0.05; Fig. 2B; Suppl. Tab. 5.2). In contrast, the experience of aggressive or passive ants did not affect the behaviour when focal ants encountered unknown non-nestmates during the test (p = 0.50; Fig. 2B; Suppl. Tab. 5.2).

**Figure 2:**
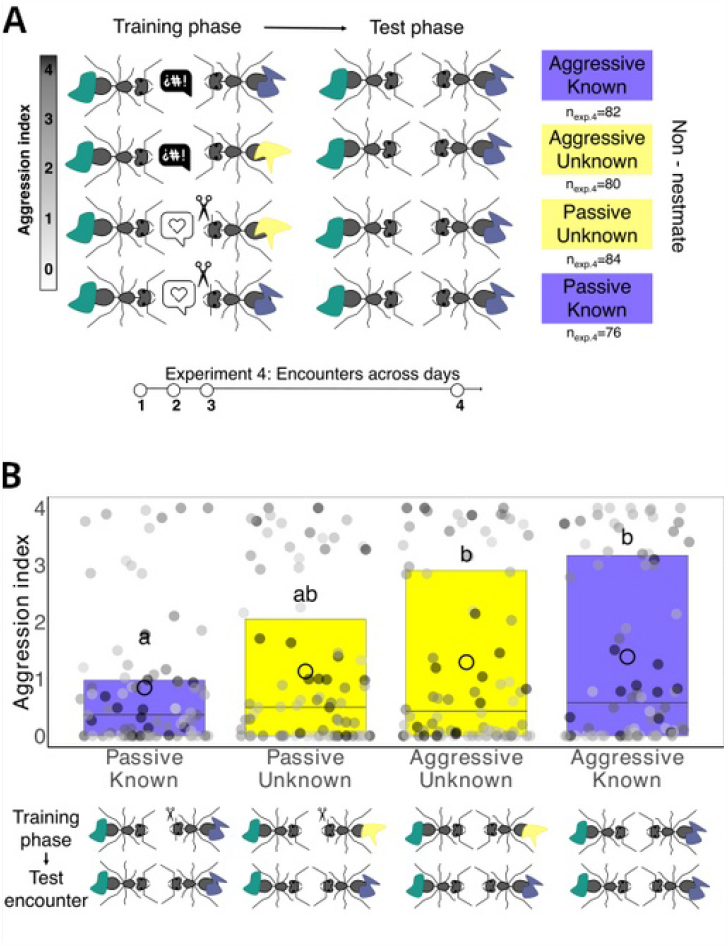
Aggression acts as the unconditioned stimulus in associative learning of non-nestmate odours. In Experiment 4 we manipulated non-nestmates into being peaceful during the training phase by cutting their antennae and encountered focal ants with either peaceful or aggressive non-nestmates once per day for three days (A). In the test phase, we encountered focal ants either with non-nestmates from the same, then known, colony, or with ants from a different, hence unknown, colony. The aggression towards known non-nestmates was lower when the previous encounters had been peaceful than if they had been aggressive (B; n = 76-84 per treatment, p < 0.05). The y-axis shows the aggression expressed by the focal ants towards non-nestmates during the test encounter. Small grey dots are measurements of individual ants’ aggression scores, the different shades correspond to the combination of focal ant and non-nestmate colony ID (3 different focal colonies). The large circles indicate means and the box plots medians and interquartile ranges. Groups with different letters were significantly different in a pairwise t-test (p < 0.05; Suppl. Tab. 5.2). The coloured shapes on the ant pictograms represent the different colony-specific odour labels. The symbols in the speech bubble indicate the tone of the encounter. <3 symbolises that the encounter was passive, !#¿ that it was agonistic.

## Discussion

The behaviour of social insect workers varies, and much of the variation is typically attributed to programmed temporal polyethism. With our experiments we demonstrated that the specific experience of individuals also contributes to the behavioural variation. We have shown that ants are more aggressive towards non-nestmates when they already had experienced individuals from the same non-nestmate colony. However, the aggression only increased if the previous encounters had been agonistic. Our results indicate that ants can associatively learn non-nestmate labels. The aggression of non-nestmates serves as the unconditioned stimulus, where the conditioned stimulus is likely to be the colony odour of the non-nestmate. Associative learning could thus allow for a more precise colony defence against intruders.

When focal ants had repeated contact with non-nestmates over multiple days, their aggression increased. This did not happen when the encounters were with nestmates, indicating that the fights that happened increased the ants’ aggression (Suppl. Fig. 3.1). However, the increase was only stable as long as the non-nestmates always came from the same colony. When we tested the ants with non-nestmates from a different and for the focal ants unknown colony, the aggression levels were as low as those of ants that had only encountered nestmates (Fig. 1B). The behavioural change is thus highly colony-specific. Most likely, the focal ants have learned the non-nestmate colony’s specific recognition labels.

When we condensed the learning and test phase into 45 minutes (experiment 3), the main pattern stayed the same but we could observe an increase of aggression also in the ants that encountered unknown non-nestmates during the test, leading to intermediate aggression levels in this treatment (Fig. 1C). This effect suggests an additional factor to play a role that increases aggression independent of the opponent’s colony. Previous studies indicated that fights alter biogenic amine and neurotransmitter levels in ants, which can lead to generally elevated aggression ^29^. Also, emotion-like states can influence decisions after learning in *Lasius niger* ants^30^. Since this effect did not occur over multiple days in our experiment 2, it is likely that the biogenic amine levels normalise over such long times.

We suspected that the ants associatively learned the recognition cues of the ants they got attacked by. In that case, the aggression an ant receives would be an unconditioned stimulus that we tried to manipulate in our fourth experiment. We modulated non-nestmate behaviour to alter the valence of the encounter and therefore the information that the focal ants could learn. Focal ants that encountered ants from a non-nestmate colony that they previously only experienced as being passive were the least aggressive of all treatments (Fig 2B). In contrast, ants were most aggressive when they had previously been attacked by these same non-nestmates. Again, the effects were colony-specific and only visible when ants encountered known non-nestmates. This highlights that the aggression an ant receives alters the valence of the recognition cues associated with it.

Focal ants being relatively peaceful after encountering passive non-nestmates is reminiscent of studies showing that ants can habituate to non-nestmate odours, leading them to treat non-nestmates as nestmates^8,13,16^. However, in contrast to these studies, our experiments exposed focal ants to non-nestmates for as little as one minute at a time. The observed effect is therefore unlikely to be due to habituation. Instead, the focal ants might have associatively learned the passive non-nestmate’s cues. This could mean that nestmate templates may also result from associative learning when ants associate the nestmate odour to a positive experience, for example receiving food or being groomed. However, we rarely observed any interaction between passive and focal ants in our training rounds. Perhaps the absence of aggression, i.e. any neutral interaction with conspecifics, might already have a positive valence.

We already know that ants can possess multiple nestmate templates^31^. For example, they might learn separate templates for foragers and nurses, which bear different CHC profiles^32,33^. Our results suggest that associative learning may play a part in generating these nestmate templates, perhaps in addition to habituation-based mechanisms^11^. We further show that the ants can learn the labels of non-nestmates that attacked them, potentially creating non-nestmate templates as well. In theory, an ant worker could collect more than just one nestmate and non-nestmate template throughout its life, and with each additional template become more efficient or precise in nestmate recognition. An alternative explanation to the ants possessing multiple templates would be that the ants possess only one nestmate template, but that this template becomes more refined when the ants learn. In the distributed model of nestmate recognition, this would mean that the border between what is perceived as a nestmate label and what is perceived as a non-nestmate label would become more accurate through experience^21^. Perceiving what corresponds to a “not-self” would allow the ants to shape their own “self” more precisely.

Independent of the exact mechanism, these processes add another mechanism causing older workers, in particular foragers, to be more aggressive: Because they spend more time outside of the nest, they have more experience with non-nestmates, which later allows them to better recognise and thus earlier attack intruders^17,34^. Since foragers will only be able to collect templates of neighbouring colonies whose foraging ranges overlap, but not of colonies further away, associative learning of non-nestmate templates can also explain the nasty neighbour effect that we found in *Lasius niger* ants from the populations studied here (experiment 1). It may, at least in part, be caused by the experience with and thus improved recognition of neighbours.

To conclude, we showed that the recognition of non-nestmates can improve through experience with these colonies. Ants can learn non-nestmate labels, most likely through associative learning with the received aggression acting as the unconditioned stimulus. Learning from non-nestmate encounters has previously been modelled^21^ but never been experimentally tested. Because individual ants will differ in the experiences they make, this process will lead to individual variation in nest defence behaviour, suggesting that there is far more individuality in ants than would be predicted by age polyethism or size polymorphism alone. On the colony level, integrating the decisions of individuals can lead to more precise colony-level decisions than individuals would ever be capable of by themselves. Associative learning is likely to play a role in the template construction all along an ant’s life, keeping the colony’s defence system up to date just like acquired immunity can protect us from novel diseases.

## Supporting information

All Supplemental Material

## Acknowledgements

We thank Patrizia d’Ettorre and Judith Korb for helpful comments on an earlier version of this manuscript and the Department of Ecology & Evolution for support during the experiments. We also thank the German Research foundation (NE 1969/6-1) for funding.

## Author contributions

CR, RE, JS, and MB conducted experiments 1-4, respectively, which had been designed by VN. MB and VN conducted the final analyses and wrote the manuscript.

## Competing interests

The authors declare no competing interests.

## Methods

### Study organism

Forty-seven colonies of *Lasius niger* were collected during spring and summer and used across the four experiments between 2016 and 2021 in Freiburg, Germany (Mooswald and Stegen). The ants were maintained at the University of Freiburg at ambient conditions in Fluon-coated plastic boxes (from 20 x 10 cm up to 20 x 20 cm according to the colony size). Each box contained a test tube filled with water and plugged with cotton. Tubes were covered with aluminium foil to create a dark nest. All colonies were provided with water ad libitum and fed with meal-worms and honey (experiments 1-2) or an artificial mixture of egg, honey agar-agar and vitamin supplement^35^ (experiments 3-4). All experiments were conducted within three months after the colonies were collected and we used a different set of colonies for each experiment.

### Experimental design

#### Experiment 2: Discrimination of known from unknown non-nestmates, long term memory

We explored how previous experiences with non-nestmates alter subsequent behaviours of focal ants. We tested whether a succession of fights with ants from a specific colony would later increase the aggression of focal ants toward that colony. During the training phase, focal ants were exposed to brief encounters with ants from a non-nestmate colony once per day over five days. The focal ants encountered a different individual from the same colony each day. As a control treatment, we set up five encounters between focal ants and their nestmates following an identical protocol. Each dyadic encounter lasted a maximum of one minute, but the ants were separated and the trial ended when the ants began to gaster-bend, to prevent a fight to death. The behaviour of the focal ants was live-recorded (see below). On day six, ants from all treatments encountered another ant for one minute. This ant could either come from the same non-nestmate colony as the non-nestmates in the training phase (“known non-nestmates”, Fig. 1A), or from another non-nestmate colony (“unknown non-nestmates”; Fig. 1A). Ants that had interacted with nestmates during the training phase also encountered non-nestmates or nestmates on day six (“nestmate treatment”, Fig. 1A; full factorial design with 9 combinations).

We used six colonies collected in 2017 and conducted 26 replicates per treatment and colony combination, for a total of 2 focal colonies x 3 training treatments (nestmates, non-nestmate colony A, non-nestmate colony B) x 3 test opponents (nestmates, non-nestmate colony A, non-nestmate colony B) x 26 replicates = 468 focal ants. The behaviour of the focal ants was live-recorded during the test.

#### Experiment 3: Discrimination of known from unknown non-nestmates, short term memory

This experiment was run analogously to experiment 2 to test whether the intervals between the training encounters affected the learning of the non-nestmate odours. Here, the training phase consisted of three short encounters of one minute each with either non-nestmates or nestmates, separated by 15 minutes intervals in a box with other nestmates. The behaviour of the focal ants was live-recorded.

After the training, we tested the aggression of all focal ants against non-nestmates on the fourth encounter. The treatments conducted were the same as in experiment 2, with the difference that no nestmate encounters took place during the test (Fig. 1A): unknown non-nestmate treatment, known non-nestmate treatment, and nestmate treatment. We used four focal colonies paired with two opponent colonies each, collected in 2022. We tested 4 focal x 2 non-nestmate colonies x 3 treatments x 3-4 replicates that amounted to 94 replicates (known non-nestmate treatment, n=33, unknown non-nestmate treatment n=30, nestmate treatment, n = 31). The behaviour of the focal ants was video-recorded.

#### Experiment 4: The role of aggression as an unconditioned stimulus

In the last experiment we tested whether the aggression an ant receives by non-nestmates acts as the unconditioned stimulus in an associative learning process. We presented focal ants with non-nestmates over three rounds of training as before, but we rendered half of the non-nestmates peaceful by ablating their antennae before the experiment. It appears that the ants become disoriented when they do not perceive olfactory input through the antennae any more (for the effect on aggression see Suppl. Fig. 5.1). In round four, we then set up encounters between focal ants that had experienced either peaceful or aggressive non-nestmates with non-nestmates from either the same or a new colony (Fig. 2A), resulting in four different combinations overall: 1) a known non-nestmate colony that was previously experienced as aggressive; 2) an unknown non-nestmate colony after previously experiencing another aggressive non-nestmate colony; 3) a known non-nestmate colony that was previously experienced as passive; 4) an unknown non-nestmate colony after previously experiencing another non-nestmate colony that was passive. We used three focal colonies paired with two opponent colonies each (all collected in 2020), for a total of 3 focal colonies x 2 non-nestmate colonies x 4 treatments x 13-15 replicates = 322 focal ants (n= 82 passive-known non-nestmates, n = 80 passive-unknown non-nestmates, n= 84 aggressive-unknown non-nestmates, and n= 76 aggressive-known non-nestmates). The behaviour of the focal ants was video-recorded.

### Behavioural aggression assays

Aggression assays were conducted in a double-circular Fluon©-coated arena placed on filter paper (Ø 5 cm). The focal ant was placed inside a smaller plastic cylinder (Ø 2.5cm) in the middle of the arena and the opponent ant was placed outside this cylinder to prevent contact. Both ants were allowed to acclimatize to the arena for five minutes before the small cylinder was removed to allow contact between the individuals and to start the trial. Aggression tests were live-or video-recorded. The focal ants’ behaviours were quantified using the softwares Etholog© version 2.2^36^ or Boris© version. 7.10.5^37^ and the observer was always blind as to the treatment and origin of the ants. The same ethogram was used across the four experiments (Suppl. Fig. 2.1). Behaviours were scored with an aggression coefficient that could take integer value from 0 to 4: no contact (a = 0), contact (a = 0), bumping (a = 1), mandible opening (a = 2), biting (a = 3), and gaster bending (a = 4). For each assay, an aggression index was calculated by multiplying the aggression coefficient by the duration (t _a_) of each behaviour expressed by the focal ant, divided by the total observation time T:

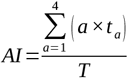

Data points were removed from the dataset for ants that were in contact for less than 1 second (17 observations in total, 7 for experiment 3 and 10 for experiment 4). We had conducted preliminary tests before experiments 2-4 to ensure that we could pair up colonies that were aggressive against each other.

### Data analysis and statistics

Across all four experiments, the aggression index (continuous variable from 0 to 4), was set up as the dependent variable in generalized linear models (GLM) with quasi-poisson error family. In experiments 2-4, the different treatments (e.g. known non-nestmates, unknown non-nestmates, nestmate treatment in experiment 2) as well as the colony origin of the non-nestmate ants were used as predictors, plus the interaction between both. Significance testing was done using variance analysis tests (anova.glm() function of the stats library). Post-hoc testing was then done using t-tests as implemented in the summary.glm() function. The results were interpreted as significant when P < 0.05 (two-tailed). All statistical analyses were performed using R 4.0.5^38^.

## Notes

### Competing Interest Statement

The authors have declared no competing interest.

## References

1. Bourke, A.F.G. Principles of Social Evolution. vol. Oxford Series in Ecology and Evolution (2011).

2. Hamilton, W. D. The genetical evolution of social behaviour. I. J. Theor. Biol. 7, 1–16, 10.1016/0022-5193(64)90038-4 (1964).

3. Hepper, P. G. Kin recognition: Functions and mechanisms a review. Biol. Rev. 61, 63–93, 10.1111/j.1469-185X.1986.tb00427.x (1986).

4. Lenoir, A., Fresneau, D., Errard, C. & Hefetz, A. Individuality and colonial identity in ants: the emergence of the social representation concept. in Information Processing in Social Insects. doi:10.1007/978-3-0348-8739-7_12. (1999)

5. Sherman, P. W, Reeve, H. K., Pfennig, D. W. Recognition systems. vol. Behavioural Ecology: An Evolutionary Approach, 4th Edition (1997).

6. van Wilgenburg, E., Felden, A., Choe, D. H., Sulc, R., Luo, J. Learning and discrimination of cuticular hydrocarbons in a social insect. Biol. Lett. 8, 17–20, 10.1098/rsbl.2011.0643 (2012).

7. Le Moli, F. & Mori, A. Field experiments on environmental sources of nestmate recognition in two species of the Formica rufa group (Hymenoptera Formicidae). Ethol. Ecol. Evol. 1, 329–339, doi.org/10.1080/08927014.1989.9525503 (1989).

8. Errard, C. & Hefetz, A. Label familiarity and discriminatory ability of ants reared in mixed groups. Insectes Soc. 44, 189–198, 10.1007/s000400050040 (1997).

9. Lenoir, A., Cuisset, D. & Hefetz, A. Effects of social isolation on hydrocarbon pattern and nestmate recognition in the ant Aphaenogaster senilis (Hymenoptera, Formicidae). Insectes Soc. 48, 101–109 101–109, 10.1007/PL00001751 (2001).

10. d’Ettorre, P., Lenoir, A. Nestmate recognition. In: Lach, L., Parr, C., Abbott, K. (Eds.), Ant Ecology. Oxford University Press, Oxford, UK, p. chapter 11, 194–208. (2010).

11. Guerrieri, F. J. Nehring, V., Jørgensen, C.G., Nielsen, J., Galizia, C.G., d’Ettorre, P. Ants recognize foes and not friends. Proc. R. Soc. B 276, 2461–2468, 10.1098/rspb.2008.1860 (2009).

12. Bos, N. & d’Ettorre, P. Recognition of Social Identity in Ants. Front. Psychology 3, 10.3389/fpsyg.2012.00083 (2012).

13. Carlin, N. F. & B. Nestmate and Kin Recognition in Interspecific Mixed Colonies of Ants. Science 222, 1027–1029, 10.1126/science.222.4627.1027 (1983).

14. Dahbi, A., Cerdá, X., Hefetz, A. & Lenoir, A. Social closure, aggressive behavior, and cuticular hydrocarbon profiles in the polydomous ant Cataglyphis iberica (hymenoptera, Formicidae). J. Chem. Ecol. 22, 2173–2186, 2173–2186, 10.1007/BF02029538 (1996).

15. Foubert, E. & Nowbahari, E. Memory span for heterospecific individuals’ odors in an ant, Cataglyphis cursor. Learning & Behavior 36, 319–326, 10.3758/LB.36.4.319 (2008).

16. Stroeymeyt, N., Guerrieri, F. J., van Zweden, J. S. & d’Ettorre, P. Rapid Decision-Making with Side-Specific Perceptual Discrimination in Ants. PLoS ONE 5, e12377 (2010).

17. Larsen, J., Nehring, V., d’Ettorre, P. & Bos, N. Task specialization influences nestmate recognition ability in ants. Behav. Ecol. Sociobiol. 70, 1433–1440, 10.1007/s00265-016-2152-9 (2016).

18. Jandt, J. & Gordon, D. The behavioral ecology of variation in social insects. Curr. Opin. Insect Sci. 15, 40–44, 10.1016/j.cois.2016.02.012 (2016).

19. Newey, P. Not one odour but two: A new model for nestmate recognition. J. Theor. Biol. 270, 7–12, 10.1016/j.jtbi.2010.10.029 (2011).

20. Nehring, V., Wyatt, T. D. & d’Ettorre, P. Noise in Chemical Communication. in Animal Communication and Noise vol. 2 373–405 10.1007/978-3-642-41494-7_13 (2013).

21. Esponda, F. & Gordon, D. M. Distributed nestmate recognition in ants. Proc. R. Soc. B Biol. Sci. 282, 20142838 (2015).

22. Temeles, E. J. The role of neighbours in territorial systems: when are they ‘dear enemies’? Anim. Behav. 47, 339–350, 10.1006/anbe.1994.1047 (1994).

23. Zorzal, G., Camarota, F., Dias, M., Vidal, D. M., Lima, E., Fregonezi, A., Campos R. I. The dear enemy effect drives conspecific aggressiveness in an Azteca-Cecropia system. Sci. Rep. 11, 6158, 10.1038/s41598-021-85070-3 (2021).

24. Langen, T. A., Tripet, F. & Nonacs, P. The red and the black: habituation and the dear-enemy phenomenon in two desert Pheidole ants. Behav. Ecol. Sociobiol. 48, 285–292, 10.1007/s002650000223 (2000).

25. Heinze, J., Foitzik, S., Hippert, A. & Hölldobler, B. Apparent Dear-enemy Phenomenon and Environment-based Recognition Cues in the Ant Leptothorax nylanderi. Ethology 102, 510–522, 10.1111/j.1439-0310.1996.tb01143.x (2010).

26. Newey, P. S., Robson, S. K. A. & Crozier, R. H. Weaver ants Oecophylla smaragdina encounter nasty neighbors rather than dear enemies. Ecology 91(8), 2366–2372, 10.1890/09-0561.1. (2010).

27. Gordon, D. M. Ants distinguish neighbors from strangers. Oecologia. 81: 198–200, 10.1007/Bf00379806 (1989)

28. Frizzi, F., Ciofi, C., Dapporto, L., Natali, C., Chelazzi, G., Turillazzi, S. G., Santini, G. The Rules of Aggression: How Genetic, Chemical and Spatial Factors Affect Intercolony Fights in a Dominant Species, the Mediterranean Acrobat Ant Crematogaster scutellaris. PLOS ONE 10, e0137919 (2015).

29. Kamhi, J. F. & Traniello, J. F. A. Biogenic Amines and Collective Organization in a Superorganism: Neuromodulation of Social Behavior in Ants. Brain. Behav. Evol. 82, 220–236, 10.1159/000356091 (2013).

30. Wenig, K., Kapfinger, H., Koch, A. & Czaczkes, T. J. Optimistic ants: Positive cognitive judgement bias but no emotional contagion in the ant Lasius niger. bioRxiv 10.1101/2022.11.03.515024. (2022)

31. Neupert, S., Hornung, M., Grenwille Millar, J. & Kleineidam, C. J. Learning Distinct Chemical Labels of Nestmates in Ants. Front. Behav. Neurosci. 12, 191, 10.3389/fnbeh.2018.00191 (2018).

32. Bonavita-Cougourdan, A., Clement, J.-L. & Lange, C. Functional subcaste discrimination (foragers and brood-tenders) in the antCamponotus vagus scop.: polymorphism of cuticular hydrocarbon patterns. J. Chem. Ecol. 19, 1461–1477, 0098-0331/93/0700-1461507.00/0 (1993).

33. Wagner, D., Tissot, M. & Gordon, D. Task-Related Environment Alters the Cuticular Hydrocarbon Composition of Harvester Ants. J Chem. Ecol., 10.1023/A:1010408725464 (2001).

34. Beshers, S. N. & Fewell, J. H. Models of division of labor in social insects. Annu. Rev. Entomol. 46, 413–440, 10.1146/annurev.ento.46.1.413 (2001).

35. Bhatkar, A. & Whitcomb, W. H. Artificial diet for rearing various species of ants. The Florida Entomologist 53, 229 (1970).

36. Ottoni, E. B. EthoLog 2.2: A tool for the transcription and timing of behavior observation sessions. Behavior Research Methods, Instruments, & Computers 32, 446–449, 10.3758/BF03200814 (2000).

37. Friard, O. & Gamba, M. BORIS : a free, versatile open‐source event‐logging software for video/audio coding and live observations. Methods Ecol. Evol. 7, 1325–1330, 1325–1330, 10.1111/2041-210X.12584 (2016).

38. R Core Team, R: A language and environment for statistical, computing, R Foundation for Statistical Computing, Vienna, Austria. URL https://www.R-project.org/. (2022).

