## Supplementary material for "Ants hold a grudge: Associative learning of non-nestmate cues improves enemy recognition": All Supplemental Material

### Supplement 1: Experiment 1, nasty neighbour effect

#### Methods

*Experimental design*

In an ecologically relevant study we investigated whether neighbouring colonies are more aggressive towards each other than distant colonies, i.e. if there is a nasty neighbour effect in *Lasius niger*. We specifically tested whether the geographic and chemical distance between colonies could predict the aggression between them. We used 24 colonies in 12 pairs collected in 2016. Both colonies of each pair came from one of two populations (Mooswald and Stegen), and were collected 1 m to 60 m apart. Within each colony pair, we set up 10 encounters between non-nestmates (12 pairs x n = 10 replicates = 120 encounters). In each encounter we recorded the behaviour of three focal ants of one colony towards one non-nestmate for two minutes (each of the two colonies was the focal colony for n = 5 replicates). Each ant was used only once. The encounters were videotaped and at any time we scored the most aggressive behaviour by one of the three ants (interactions ranged from antennation to gaster bending, Suppl. Fig. 2.1). We similarly staged five additional encounters between nestmates of each colony (24 colonies x 5 replicates = 120 encounters). We analysed the cuticular chemicals of n = 3-6 ants per colony (113 samples in total) and used the data to calculate the chemical distance between the colonies to see if it would explain the aggression between the colonies.

*CHC analysis*

To assess the effect of chemical differences between colonies on the aggression between colonies, we quantified the cuticular hydrocarbon profiles of 3-4 ants per colony. After we killed the individuals by freezing, CHCs were extracted by immersing single ants in 200 µL pentane for three minutes. The pentane was allowed to evaporate and the extracts were stored at -20°C until further analysis. The samples were re-diluted in 15 µL heptane, 3 µL of which were injected into a gas chromatograph coupled with a mass spectrometer (GC-MS). The GC (7890B, Agilent Technologies, Germany) was equipped with a HP-5MS Ultra Inert 5% Phenyl-Methylpolysiloxane column (0.25µm, 30m, 0.25mm, Agilent Technologies). The inlet was operated in splitless mode at 250°C. The flow rate was set to 1 mL/min and helium was used as the carrier gas. The column temperature was increased from 70°C to 220°C by 10°C/min, then from 220°C to 320°C by 5°C/min, and was held constant for another 4 minutes. GC peaks were transferred to a mass selective detector (5977A, Agilent Technologies, Germany) and fragmented by a ionization voltage of 70eV. Fragments in the range of 50 – 700 m/z were detected and used for substance quantification and identification. For identification, we used retention indices and diagnostic ions (Eberlin 2006). We quantified the areas of all hydrocarbon peaks in MSD ChemStation (F.01.01.2317, Agilent Technologies).

We calculated the average hydrocarbon profile per colony (colony centroids) as the average relative peak area per substance. The Bray Curtis distance between colony centroids (package “vegan”, Oksanen et al. 2022) served as a measure of chemical dissimilarity.

*Data analysis and statistics*

The aggression index (Suppl. Fig. 2.1) was set up as the dependent variable in generalized linear models (GLM) with quasi-poisson error family. The geographic and chemical distances between colonies were used as predictors. Since the experiment was conducted in two populations, the population ID and its interaction with distance were also included. P-values were calculated by variance analysis (ANOVA function of stats library) and Wald tests (summary function of stats library). The results were interpreted as significant when $P < 0.05$ (two-tailed). All statistical analyses were performed using R 4.0.5 software (R Core Team, 2022).

### Results

Aggression was higher between non-nestmates than among nestmates (Suppl. Fig. 1.1). Non-nestmate aggression was lowest among colonies of medium distance (Suppl. Fig. 1.2) and best described by a unimodal model (n = 120, quasibinomial glm with distance $F_{1,119} = 6.2$, $p = 0.01$ distance$^2$ $F_{1,117} = 60.79$, $p < 0.001$), with no big differences between the two populations (population $F_{1,116} = 1.4$, $p = 0.24$; distance : population $F_{1,115} = 0.51$, $p = 0.48$; distance$^2$ : population $F_{1,114} = 0.02$, $p = 0.88$). Since we suspected that the elevated aggression among adjacent colonies resulted from the nasty neighbour effect caused by repeated interactions between colonies, we correlated aggression with distance again, but this time excluded neighbouring colonies. According to our own estimates and data from the literature, *Lasius niger* forage up to ca. 3 m (Headley 1941, Devigne & Detrain, 2006). We thus excluded colony pairs that were less than three meters apart, resulting in a dataset containing only colony pairs that were 8 m or further apart. Then, the quadratic distance term became less important (n = 90; distance $F_{1,88} = 55.7$, $p < 0.001$; distance$^2$ $F_{1,87} = 5.5$, $p < 0.05$; population $F_{1,86} = 0.9$, $p = 0.35$; distance : population $F_{1,85} = 2.5$, $p = 0.11$; distance$^2$ : population $F_{1,84} = 0.8$, $p = 0.37$).

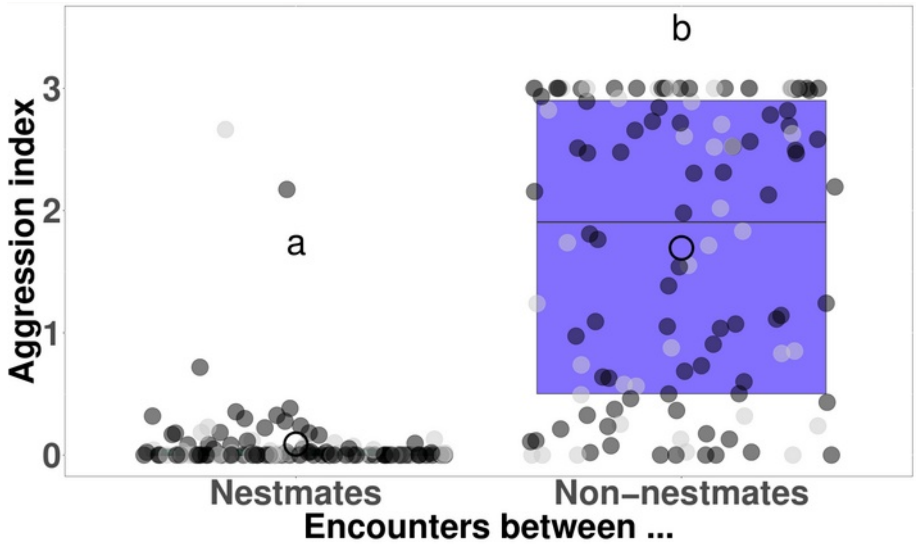


**Supplementary Figure 1.1:** *Lasius niger* workers were more aggressive towards non-nestmate than towards nestmate workers (n = 240; quasipoisson glm nestmate $F_{1,238} = 215.6$, $p < 0.001$; population $F_{1,237} = 0.0$, $p > 0.99$; interaction nestmate : population $F_{1,236} = 0.0$, $p = 0.88$). The large circle indicates the mean, the boxplots indicate median and first and third quartiles. Dots are measurements of individual encounters, with colours indicating the population identity.

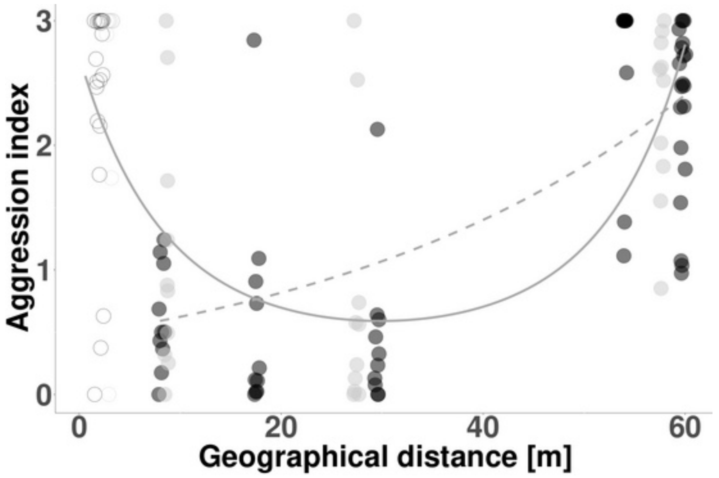


**Supplementary Figure 1.2:** Nasty neighbour effect in *Lasius niger.* Ants from colonies within foraging distance of each other (circles) are more aggressive than colonies that are further apart (filled dots). The overall pattern follows a parabola (solid line; prediction from a quasipoisson model with distance$^2$ as a predictor) but aggression among non-neighbours increases with distance (dashed line, regression without quadratic term). Grey dots are measurements of individual ants' aggression score, the different shades of grey correspond to the two populations. Darker shades indicate overlapping data points.

Chemical distance between colonies positively affected the aggression between non-neighbouring colonies in both populations (Suppl. Fig. 1.3; n = 90: quasibinomial glm with population $F_{1,88} = 0.1$, $p = 0.72$; chemical distance $F_{1,87} = 34.2$, $p < 0.001$; population : chemical distance $F_{1,86} = 0.7$, $p = 0.4$). An increase in chemical distance can explain why colonies become more aggressive when they are further apart.

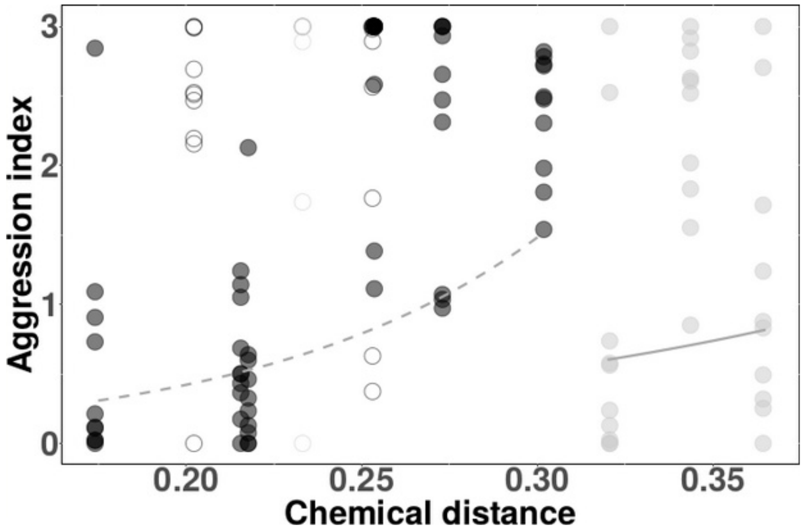


**Supplementary Figure 1.3:** Aggression increased with chemical distance between non-neighbouring colonies (filled circles) in both populations (dark grey, dashed line). The chemical distances and their variation were different between the two populations (dark grey dots, dashed line; light grey dots, solid line). The lines are quasipoisson regressions for non-neighbouring colony pairs (filled circles), while neighbouring colony pairs are depicted with open circles.

#### Discussion

Neighbouring colonies within foraging distance were particularly aggressive towards each other, and this effect was restricted to the assumed foraging distance for this species (ca. 5 m; Pontin 1963, Headley 1941, Devigne & Detrain, 2006). Colony pairs 8 m – 50 m apart were relatively peaceful. We did observe aggression to increase again between colonies that were more than 50 m apart, which may be caused by these colonies differing strongly in their recognition cues. Our analysis of the cuticular hydrocarbons is consistent with this assumption because aggression of nopn-neighbours increased with chemical distance. When labels (cuticular hydrocarbon profiles) are too different, ants are known to react aggressively (Dahbi et al. 1996), which might explain why some studies seemed to find a combination of the nasty neighbour and dear enemy effects.

#### References

Eberlin, M. N. Structurally diagnostic ion/molecule reactions: class and functional-group identification by mass spectrometry. *J. Mass Spectrom.* **41**, 141–156, 141–156, 10.1002/jms.998 (2006).

Oksanen, J. *et al.* vegan: Community Ecology Package. (2011).

R Core Team, R: A language and environment for statistical, computing, R Foundation for Statistical Computing, Vienna, Austria. URL https://www.R-project.org/. (2022).

Headley, A. E. A Study of Nest and Nesting Habits of the Ant *Lasius niger* Subsp. *alienus* Var. *americanus* Emery. *Annals of the Entomological Society of America* **34**, 649–657, 10.1093/aesa/34.3.649 (1941).

Devigne, C. & Detrain, C. How does food distance influence foraging in the ant *Lasius niger*: the importance of home-range marking. *Insectes Sociaux* **53**, 46–55 (2006).

Pontin, A. J. Further Considerations of Competition and the Ecology of the Ants *Lasius flavus* (F.) and *L. niger* (L.). *The Journal of Animal Ecology* **32**, 565 (1963).

Dahbi, A., Cerdá, X., Hefetz, A. & Lenoir, A. Social closure, aggressive behavior, and cuticular hydrocarbon profiles in the polydomous ant *Cataglyphis iberica* (hymenoptera, Formicidae). *J. Chem. Ecol.* **22**, 2173–2186, 10.1007/BF02029538 (1996).

### Supplement 2: Methods

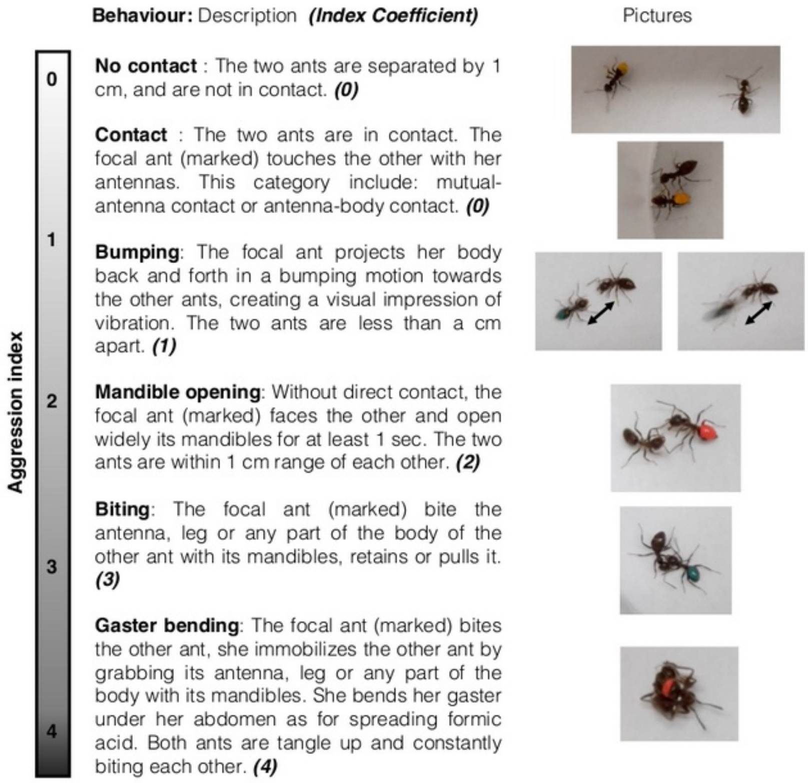


**Supplementary Figure 2.1:** Ethogram used across all experiments.

### Supplement 3: Experiment 2, learning over 5 days

A

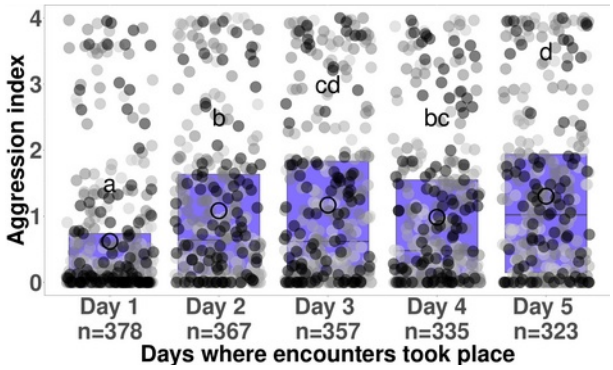


B

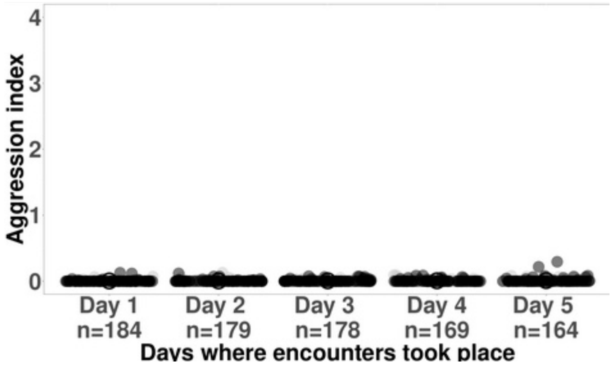


**Supplementary Figure 3.1: Training phase.** The aggression of *Lasius niger* workers during the training phase significantly increased through five consecutive days of encounters with non-nestmates (A; n = 1760 encounters; quasipoisson GLM days $F4_{,1752} = 18.84$, $p < 0.001$, opponent colony origin $F_{3,1756} = 26.69$, $p < 0.001$; opponent colony origin : days $F_{12,1740} = 0.87$, $p = 0.58$), but not with nestmates (B; n = 874 encounters; quasipoisson GLM days $F_{4,868} = 2.0$, $p = 0.09$; opponent colony origin: $F_{1,872} = 1.2$, $p = 0.27$; interaction opponent colony origin : days: $F_{4,864} = 0.5$, $p = 0.70$). Small grey dots are measurements of individual ants' aggression score, the different shade correspond to the colony the focal ants belonged to. The large circle indicates the mean, the boxplots indicate median and first and third quartiles. Groups with different letters were significantly different in pairwise t-tests ($p < 0.05$).

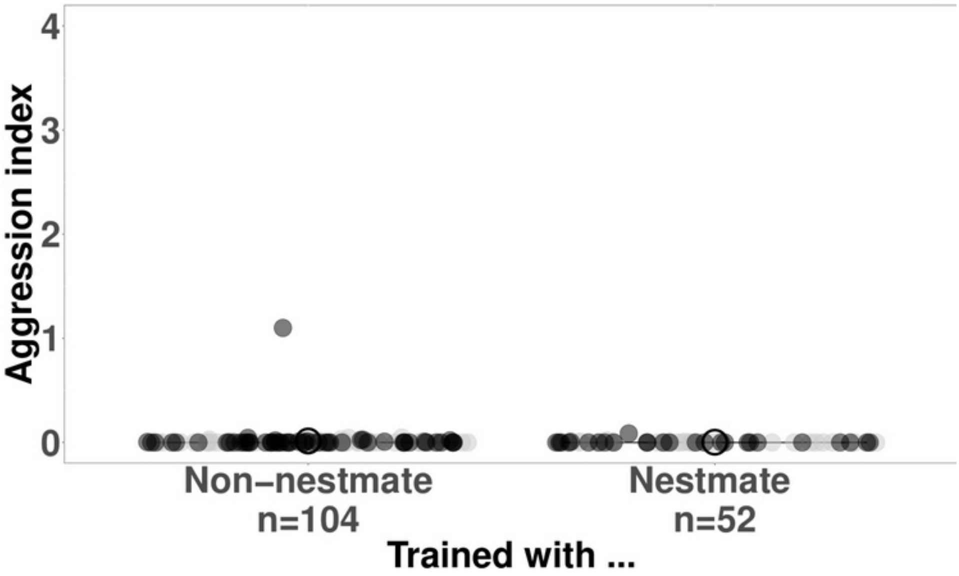


**Supplementary Figure 3.2:** Focal ants were generally peaceful towards their nestmates during the test. Their aggression was not affected by whether they had been trained with nestmates or non-nestmates. Small grey dots are measurements of individual ants' aggression score, the different shades correspond to the focal ants' colony IDs. The large circle indicates the mean, box plots are not visible because first and third quartiles were zero in both treatments.

**Supplementary Table 3.1:** Analysis of Variance on a GLM with quasi-poisson error family, for ants that encountered non-nestmates on day 6: Full model: AI ~ Opponent colony origin x Treatment

Opponent colony origin: $F_{3,308} = 2.3$, $p < 0.001$
Treatment: $F_{2,306} = 13.84$, $p < 0.001$
Opponent colony origin : Treatment: $F_{6,300} = 1.53$, $p = 0.17$

**Supplementary Table 3.2:** Pairwise comparisons between treatments, p-values from t-tests, for ants that encountered non-nestmates on day 6: Full model: AI ~ Opponent colony origin + Treatment

known non-nestmates vs. nestmates: $t = -4.2$, $p < 0.001$
known non-nestmates vs. unknown non-nestmates: $t = -4.5$, $p < 0.001$
nestmates vs. unknown nestmates $t = -0.3$, $p = 0.74$

**Supplementary Table 3.3:** Analysis of Variance on a GLM with quasi-poisson error family, for ants that encountered nestmates on day 6. Full model: AI ~ Opponent colony origin x Treatment

Full model: AI ~ Opponent colony origin x Training treatment
Opponent colony origin: $F_{1,154} = 2.4305$, $p = 0.1211$
Treatment: $F_{1,153} = 1.6903$ $p = 0.1955$
Opponent colony origin : Training treatment: $F_{4,864} = 0.5457$, $p = 0.70219$

### Supplement 4: Experiment 3, learning over 45 minutes

A B

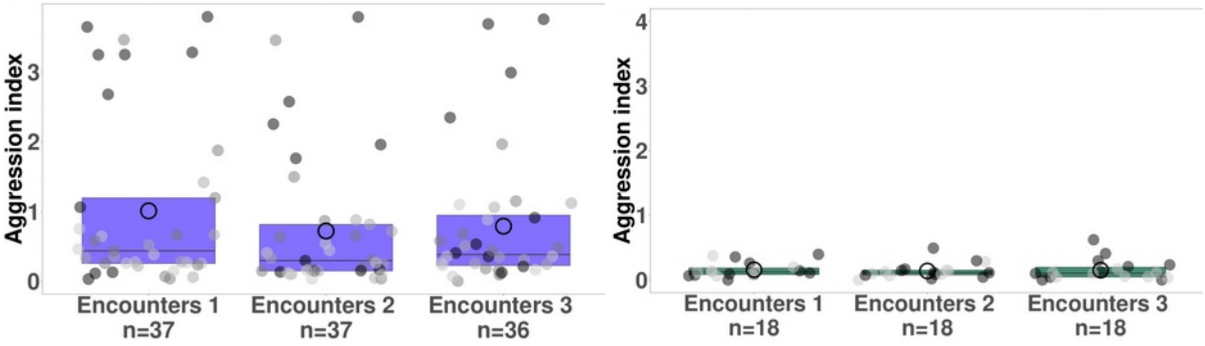


**Supplementary Figure 4.1:** During three training encounters with non-nestmates (A), the aggression of focal ants did not significantly increase (n = 110; quasibinomial GLM $F_{2,104} = 1.3$, p= 0.28). The combination of colonies affected the aggression ($F_{3,106}$= 14.4, p < 0.001), but there was no interaction with the number of the training encounter ($F_{6,98} = 0.3$, p = 0.27). During the encounters with their nestmates, focal ants were typically peaceful, independent of the number of encounters and colony (colony origin: $F_{1,52} = 2.5$, p = 0.12; encounter $F_{2,50} = 0.1$, p = 0.91; interaction colony : encounter $F_{2,48}$ = 1.1, p = 0.33). Small grey dots are measurements of individual ants, the different shades correspond to their colony IDs. The large circle indicates the mean, and boxplots indicate medians and interquartile ranges. Groups with different letters were significantly different in a t-test (p < 0.05). Green represents the group of ants that have encountered nestmate during the training phase, purple represents the group of ants that encountered non-nestmates.

**Supplementary Table 4.1:** GLM with quasi-poisson error family, p value calculated by ANOVA, F-test. Full model: AI ~ Opponent colony origin x Treatment

Opponent colony origin: $F_{5,89}$ = 6.20, p < 0.001***
Treatment: $F_{2,87}$ = 4.7915, p = 0.01093 *
Opponent colony origin : Treatment: $F_{10,77}$ = 1.6371, p = 0.11190

**Supplementary Table 4.2:** Pairwise comparisons between treatments, for the model AI ~ Opponent colony origin + Treatment

known non-nestmates vs. nestmates: t = -2.572, p = 0.01179 *
known non-nestmates vs. unknown non-nestmates: t = -1.928, p = 0.05712 .
nestmates vs. unknown nestmates t = 0.797, p = 0.42785

### Supplement 5: Experiment 4, aggression as an unconditioned stimulus

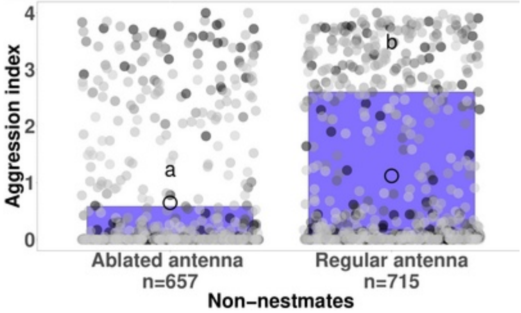


**Supplementary Figure 5.1:** *Lasius niger* workers significantly decreased their aggression towards non-nestmates when their own antennae had been ablated (n = 1372; quasiposson GLM, $F_{1,1365} = 48.9$, $p < 0.001$). The combination of colonies also affected aggression (GLM, $F_{5,1366} = 7.0$, $p < 0.001$), but this effect was restricted to the ants that still had intact antennae (interaction colony origin : antenna treatment $F_{5,1360} = 2.5$, $p < 0.05$). Data presented in this figure stem from the training phase of the experiment (Suppl. Fig. 5.2) and show the aggression expressed by ants with or without antennae towards non-nestmates. Small grey dots are measurements of individual ants' aggression scores, the different shades correspond to their colony IDs. The large circles indicate the mean, and boxplots indicate medians and interquartile ranges. Groups with different letters were significantly different in a t-test ($p < 0.05$).

A

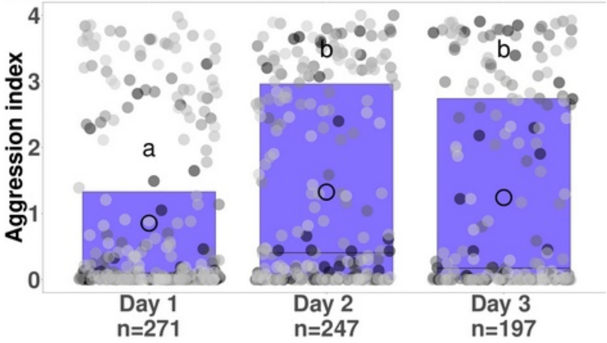


B

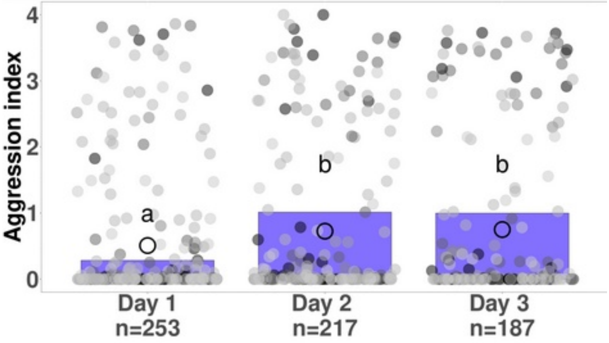


**Supplementary Figure 5.2:** The aggression of *Lasius niger* workers during the training phase significantly increased through three consecutive days of encounters towards aggressive non-nestmates (A; n = 657; quasipoisson GLM: $F_{2,707} = 8.4$, $p < 0.001$). The combination of colonies also affected aggression ($F_{5,709} = 5.1$, $p < 0.001$), but there was no interaction between colony origin and the day of the encounter (origin : days: $F_{10,697} = 0.8$, $p = 0.58$). Aggression also increased against non-nestmates that were passive because we had cut off their antennae, but to a lesser extent (B; n = 657; colony origin: $F_{5,651} = 5.0$, $p < 0.001$; factor days: $F_{2,649} = 3.6$, $p < 0.05$; interaction origin : days $F_{10,639} = 0.6$, $p = 0.84$). Small grey dots are measurements of individual aggression scores, the different shades correspond to the ants' colony IDs. The large circle indicates the mean, and boxplots indicate medians and interquartile ranges. Groups with different letters were significantly different in a t-test ($p < 0.05$).

**Supplementary Table 5.1:** Analysis of Variance on a GLM with quasi-poisson error family.
Full model: AI ~ Opponent colony origin x Treatment

Opponent colony origin: $F_{5,320}$ = 3.4986, p= 0.004334 **
Treatment: $F_{3,317}$ = 2.9656, p = 0.032322 *
Opponent colony origin : Treatment: $F_{15,302}$ = 0.7149, p = 0.769248

**Supplementary Table 5.2:** Pairwise comparisons between treatments (t-test):
Full model: AI ~ Opponent colony origin + Treatment

Aggressive known non-nestmates vs. passive known non-nestmates: t = -2.407, p = 0.01666 *
Aggressive known non-nestmates vs. passive unknown non-nestmates: t = -1.027 , p = 0.30504
Aggressive known non-nestmates vs. aggressive unknown non-nestmates: t = -0.366, p = 0.71495
Passive known non-nestmates vs. passive unknown non-nestmates: t = 1.417 , p = 0.15747
Passive known non-nestmates vs. aggressive unknown non-nestmates: t = 2.085, p = 0.03786 *
Passive unknown non-nestmates vs. aggressive unknown non-nestmates: t = 0.677, p = 0.49892
